# CBP80/20-dependent translation initiation factor (CTIF) inhibits HIV-1 Gag synthesis by targeting the function of the viral protein Rev

**DOI:** 10.1101/710137

**Authors:** Francisco García-de-Gracia, Daniela Toro-Ascuy, Sebastián Riquelme-Barrios, Camila Pereira-Montecinos, Bárbara Rojas-Araya, Aracelly Gaete-Argel, Mónica L. Acevedo, Jonás Chnaiderman, Fernando Valiente-Echeverría, Ricardo Soto-Rifo

## Abstract

Translation initiation of the human immunodeficiency virus type-1 (HIV-1) unspliced mRNA has been shown to occur through cap-dependent and IRES-driven mechanisms. Previous studies suggested that the nuclear cap-binding complex (CBC) rather than eIF4E drives cap-dependent translation of the unspliced mRNA and we have recently reported that the CBC subunit CBP80 supports the function of the viral protein Rev during nuclear export and translation of this viral transcript. Ribosome recruitment during CBC-dependent translation of cellular mRNAs relies on the activity CBP80/20 translation initiation factor (CTIF), which bridges CBP80 and the 40S ribosomal subunit through interactions with eIF3g. Here, we report that CTIF restricts HIV-1 replication by interfering with Gag synthesis from the unspliced mRNA. Our results indicate that CTIF associates with Rev through its N-terminal domain and is recruited onto the unspliced mRNA ribonucleoprotein complex in order to block translation. We also demonstrate that CTIF induces the cytoplasmic accumulation of Rev impeding the association of the viral protein with CBP80. We finally show that CTIF restricts HIV-2 but not MLV Gag synthesis indicating an inhibitory mechanism conserved in Rev-expressing human lentiviruses.

## INTRODUCTION

Ribosome recruitment onto the HIV-1 unspliced mRNA has been shown to occur through cap-dependent and cap-independent mechanisms (1-3). Initial studies showed that the highly structured 5’-untranslated region (5’-UTR) present within the full-length unspliced mRNA was inhibitory for translation in cell-free translation systems suggesting that cap-dependent ribosome scanning was not an efficient mechanism of translation initiation operating in HIV-1 transcripts (4-6). In agreement with this idea, several groups have reported that the 5’-UTR of the HIV-1 unspliced mRNA harbors a cell cycle-dependent internal ribosome entry site (7-11). However, more recent reports including ours have shown that cap-dependent ribosome scanning operates on the unspliced mRNA in a mechanism supported by the host DEAD-box RNA helicase DDX3 but independent of the cap-binding protein eIF4E (12-16). In this sense, HIV-1 unspliced mRNA translation was also proposed to occur in a nuclear cap-binding complex (CBC)-dependent fashion during the virally induced arrest of the cell cycle (17). Consistent with these observations, we have recently reported that the CBC subunit CBP80 (NCBP1) promotes nuclear export and translation of the HIV-1 unspliced mRNA by supporting the functions of the viral protein Rev on these processes (18).

The CBC is recruited onto the nascent transcript early during transcription and has been involved in every step of mRNA metabolism from transcription to splicing, nuclear export, translation and decay (19). Two different CBCs have been described so far in mammalian cells, the canonical CBC composed by CBP20 (NCBP2) and CBP80 and a recently described CBC in which CBP80 interacts with the newly discovered cap-binding protein NCBP3 (20,21). CBC-dependent translation has mostly been associated to the mRNA quality control mechanism known as nonsense-mediated mRNA decay (NMD), which triggers the degradation of mRNA harboring premature stops codons (22-24).

Translation initiation driven by the CBC relies on the CBP20/80 translation initiation factor (CTIF), which binds to the CBP80 subunit and interacts with eIF3 to promote 40S small ribosomal subunit recruitment (25,26). CTIF also recruits the DEAD-box RNA helicase eIF4AIII, which drives the unwinding of RNA structures on the 5’-UTR of mRNA translated through a CBC-dependent mechanism (27). More recently, it was shown that CTIF couples translation with autophagy by driving misfolded poplypetides to the aggresomes (28). Here, we report that CTIF is a potent inhibitor of Gag synthesis that blocks unspliced mRNA translation.

We observed that CTIF interacts with Rev through its N-terminal and is incorporated into the unspliced mRNA ribonucleoprotein complex. We also demonstrate that CTIF induces the cytoplasmic accumulation of Rev and interferes with the association between the viral protein and CBP80. We finally shown that CTIF interferes with Gag synthesis from HIV-2 but not MLV, indicating a conserved mechanism by which CTIF targets the function of human lentiviral Rev proteins on Gag synthesis.

Together, our data provide evidence for a novel restriction of HIV-1 replication, which is exerted at the level of unspliced mRNA translation and targets the function of the viral protein Rev.

## MATERIALS AND METHODS

### DNA constructs

The pNL4.3-ΔEnv, pNL4.3R, pNL4.3R-ΔRev, pROD10R and pNCAC proviral vectors were previously described (18,29,30). The pcDNA3-Flag-CTIF, pcDNA3-Flag-CTIF(1-305) and pcDNA3-Flag-CTIF(306-598) were previously described (25). The pcDNA-d2EGFP, pCDNA β-globin 5’-UTR, pCIneo-Renilla and pEGFP-Rev were previously described (18,29).

### Cell culture and DNA transfection

HeLa and HEK293T cells were maintained with DMEM (Life Technologies) supplemented with 10% FBS (Pan-Biotech) at 37 °C and a 5% CO2 atmosphere. The Jurkat clone E6-1 was maintained in RPMI 1640 (Life Technologies) supplemented with 10% FBS (Pan-Biotech) and antibiotics (Sigma-Aldrich) at 37 °C and a 5% CO2 atmosphere. Cells were transfected using linear PEI ∼25,000 Da (Polysciences) as described previously (18,29).

### Analysis of Renilla and firefly luciferase activities

Renilla luciferase activity was determined using the Renilla Reporter Assay System (Promega) and Renilla/firefly luciferase activities were determined using the Dual-Luciferase® Reporter Assay System (Promega) in a GloMax® 96 microplate luminometer (Promega).

### Western blot

Cells extract were subjected to 10% SDS-PAGE and transferred to an Amersham Hybond™-P membrane (GE Healthcare). Membranes were incubated with an HIV-1 p24 monoclonal antibody diluted to 1/1000 (Catalog number 3537), a rabbit anti-CTIF antibody (ThermoFisher Scientific) diluted to 1/250, a mouse anti-GAPDH antibody (Santa Cruz Biotechnologies) diluted to 1/5000, a rabbit anti-MLV CA (kindly provided by Dr. Gloria Arriagada, UNAB, Chile) diluted 1/1000, a mouse anti-F protein of HRSV (Santa Cruz Biotechnologies) diluted 1/500, or mouse anti-puromycin antibody (Milipore, clone 12D10) and anti-actin antibody (Santa Cruz Biotechnologies) diluted 1/750. Upon incubation with the corresponding HRP-conjugated secondary antibody (Jackson ImmunoResearch) diluted to 1/5000. Membranes were revealed with the Pierce® ECL substrate or SuperSignal™ West femto (Thermo Scientific) using a Mini HD9 Western blot Imaging System (UVItec).

### CTIF knock down, stable cell line generation, proviral transfection and recovery assay

Lentiviral particles carrying an anti-CTIF shRNA were produced in HEK293 cells by transfecting a commercially available pLKO.1 vector containing the shRNA sequence targeting the 3’-UTR of the CTIF mRNA (Sigma-Aldrich), pVSVg and psPax2. Supernatants were collected at 48 hours post transfection, cleared through a 0.22 µm filter and used to transduce HeLa cells for 24 hours. The medium containing the lentiviral particles was replaced by fresh DMEM and cells were grown for additional 48 hours. Finally, cells were treated with puromycin (10 μg/mL) for 10 days at 37°C and 5% CO_2_. CTIF knockdown was evaluated by Western blot as mentioned above. Lentiviral particles carrying a scramble shRNA were used as a control. The sequences of CTIF and scramble shRNAs are in Supplementary Table 1.

Control and CTIF knockdown cells were transfected with pNL4.3R together with pcDNA-d2EGFP or pcDNA3-Flag CTIF, and Renilla luciferase activity was determinated as mentioned above.

### Pseudotyped virus production and infection assays

HEK293T cells were co-transfected with the pNL4.3-ΔEnv provirus and pVSVg and cell supernatants were collected at 72 hour post-transfection, cleared through a 0.22 µm filter and used to infect Jurkat cells by using 1 volume virus stocks per volume of cells. Infected cells were recovered at 8, 12, 20, 24, 48, 72, 96 hours post-infection and washed with 300 μl of PBS and pelleted at 500 x g for 5 min at 4 °C. Cells were resuspended in 100 µl of ice cooled lysis Buffer I (150 mM sodium chloride, 1.0% NP-40, 0.5% sodium deoxycholate, 50 mM Tris-HCl, pH 7.5 and protease inhibitor [Roche]), vortexed for 5 seconds, incubated for 30 min at 4°C under agitation and centrifuged at 12,000 rpm for 20 min at 4°C. The supernatant containing the whole cellular lysate was recovered and 30 µg of total protein were used for Western blot analysis. For hRSV production HEp-2 cells were infected with 500 µl of the virus and supernantants were recovered when an 80% of cytopathic effect was observed. Supernatants were cleared by centrifugation and stored at −80°C. For quantification of hRSV we perform a TCID_50_ in the MA104 cell line as previously described (31).

### Surface Sensing of Translation (SUnSET) assay

Global proteins synthesis under control or CTIF overexpression was analyzed using the puromycin incorporation into new peptide synthesis as previously described (32). Briefly, HeLa cells were transfected with pcDNA-d2EGFP or pcDNA3-Flag-CTIF and treated with 10 µg/ml of puromicine for 10 min at 37°C and 5% CO_2_ at 24 hpt. Cells were washed with PBS twice and lysed with lysis Buffer II (10 mM Tris-HCl pH 7.5, 100 mM NaCl, 0.5% NP-40, 1 mM EDTA and EDTA-free protease inhibitors cocktail [Roche]). As a control of protein synthesis inhibition, we treated HeLa cells with 0.5 mM of arsenite (Sigma-Aldrich) for 45 min prior to puromycin treatment. 20 µg of total proteins were loading in a SDS gel for Western blot analysis.

### RNA extraction and RT-qPCR

Cytoplasmic RNA extraction and RT-qPCR from cytoplasmic RNA were performed exactly as we have previously described (18,29). The GAPDH housekeeping gene was amplified in parallel to serve as a control reference. Relative copy numbers of HIV-1 unspliced mRNA were compared to GAPDH using x^−ΔCt^ (where x correspond to the experimentally calculated amplification efficiency of each primer couple). Sequences of the primers used and the experimentally calculated amplification efficiency of the primers are presented in Supplementary Table 2.

### RNA fluorescent *in situ* hybridization, immunofluorescence and confocal microscopy

RNA FISH and immunofluorescence analyses were performed exactly as we have recently reported but using an anti-Flag (Sigma-Aldrich) instead of an anti-HA primary antibody (18).

### Proximity ligation assay (PLA) and *in situ* hybridization coupled to PLA (ISH-PLA)

PLA and ISH-PLA were carried out using mouse anti-digoxin (Roche) and rabbit anti-Flag (Sigma-Aldrich), the DUOLINK II In Situ kit (Sigma-Aldrich) and PLA probe anti-mouse minus and PLA probe anti-rabbit plus (Sigma-Aldrich) exactly as we have recently described (18).

## RESULTS

### CTIF regulates HIV-1 Gag synthesis during viral replication

We have recently reported that CBP80 promotes Gag synthesis from the unspliced mRNA in association with the viral protein Rev (18). Since CTIF bridges CBP80 and the 40S ribosomal subunit by interacting with eIF3g during CBC-dependent translation, we sought whether CTIF was involved in gene expression from the HIV-1 unspliced mRNA. We first analyzed the expression of the endogenous protein in HIV-1-infected T-cells and observed that the presence of HIV-1 induced an increase in the levels of CTIF from 8 to 24 hour post-infection to then decrease at later time points (Fig. 1A, upper panel). We consistently observed that the drop in the levels of CTIF was correlated with the accumulation of the Gag polyprotein (Figure 1A, lower panel), which led us to speculate that increased levels of CTIF are not required for the synthesis of Gag. Consistent with such hypothesis, overexpression of Flag-tagged CTIF in HeLa cells resulted in a strong inhibition of HIV-1 Gag synthesis from the pNL4.3R reporter provirus (Fig. 1B). It should be mentioned that Flag-CTIF overexpression had no impact on gene expression from a reporter vector, on global protein synthesis or in other viral proteins such as Vif suggesting a specific effect on HIV-1 Gag synthesis (Supplementary Figs. 1A, 1B and 1C). In agreement with a negative effect of CTIF on Gag synthesis, we observed that even a mild reduction of the endogenous levels of CTIF resulted in enhanced levels of Gag synthesis, which were further reduced when Flag-CTIF was ectopically expressed (Fig. 1C).

**Figure 1:**
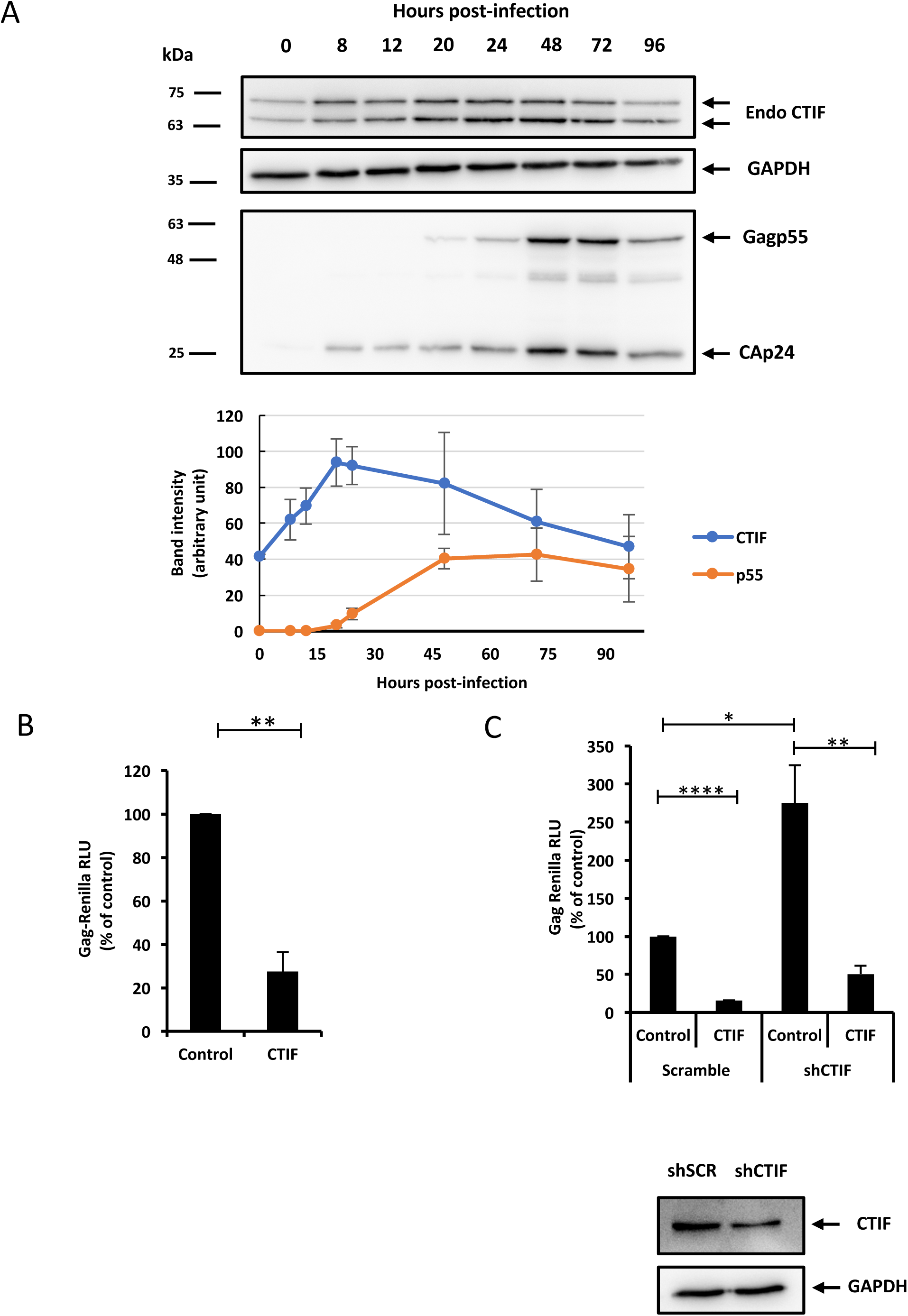
CTIF inhibits Gag synthesis from the HIV-1 unspliced mRNA. A) Jurkat cells were infected with VSVg-pseudotyped HIV-1 and cell extracts prepared at 0, 8, 12, 20, 24, 48, 72 and 96 hours post-infection were used to detect Gag (and its processing intermediates) and CTIF by Western blot. GAPDH was used as loading control. Band intensity for CTIF and Gag (pr55) from three independent experiments were quantified, normalized to GAPDH and plotted (n = 3, +/- SEM). B) HeLa cells were transfected with 1 µg of pcDNA-d2EGFP (Control) or pcDNA3-Flag-CTIF together with 0.3 µg of pNL4.3R as described in materials and methods and Renilla activity was determined 24 hpt. Results were normalized to the control (arbitrary set to 100%) and correspond to the mean +/- SD of three independent experiments (**P < 0.01, t-test). In parallel, cells extracts were used to detect Flag-CTIF by Western blot for expression control. Actin were used as a loading control. C) Scramble and CTIF knockdown HeLa cells were transfected with 1 µg of pcDNA-d2EGFP (used as a control) or pcDNA3-Flag-CTIF together with 0.3 µg of pNL4.3R as described in materials and methods. Renilla luciferase activity was determined at 24 hpt. Results were normalized to the control (arbitrary set to 100%) and correspond to the mean +/- SD of three independent experiments (*P < 0.05; **P < 0.01 and ****P < 0.0001, t-test). Extracts from HeLa Scramble and CTIF knockdown cells were used to verify CTIF knockdown by Western blot. GAPDH was used as a loading control.

Taken together these results indicate that CTIF is a negative regulator of Gag synthesis during HIV-1 replication.

### CTIF inhibits Gag synthesis in a Rev-dependent manner

CTIF is a CBC-dependent translation initiation factor that is recruited to the mRNP upon nuclear export and was shown to play a scaffold role during the pioneer round of translation (25,26). Thus, in order to identify the step at which CTIF was interfering with Gag synthesis, we analyzed the impact of CTIF overexpression on the cytoplasmic levels of the unspliced mRNA. Despite we observed a strong inhibition of Gag synthesis under ectopic expression of CTIF, the cytoplasmic levels of the unspliced mRNA levels were minimally affected. However, this reduction in the cytoplasmic unspliced mRNA does not explain the strong decrease in Gag indicating that CTIF might exert its effects on translation (Fig. 2A).

**Figure 2:**
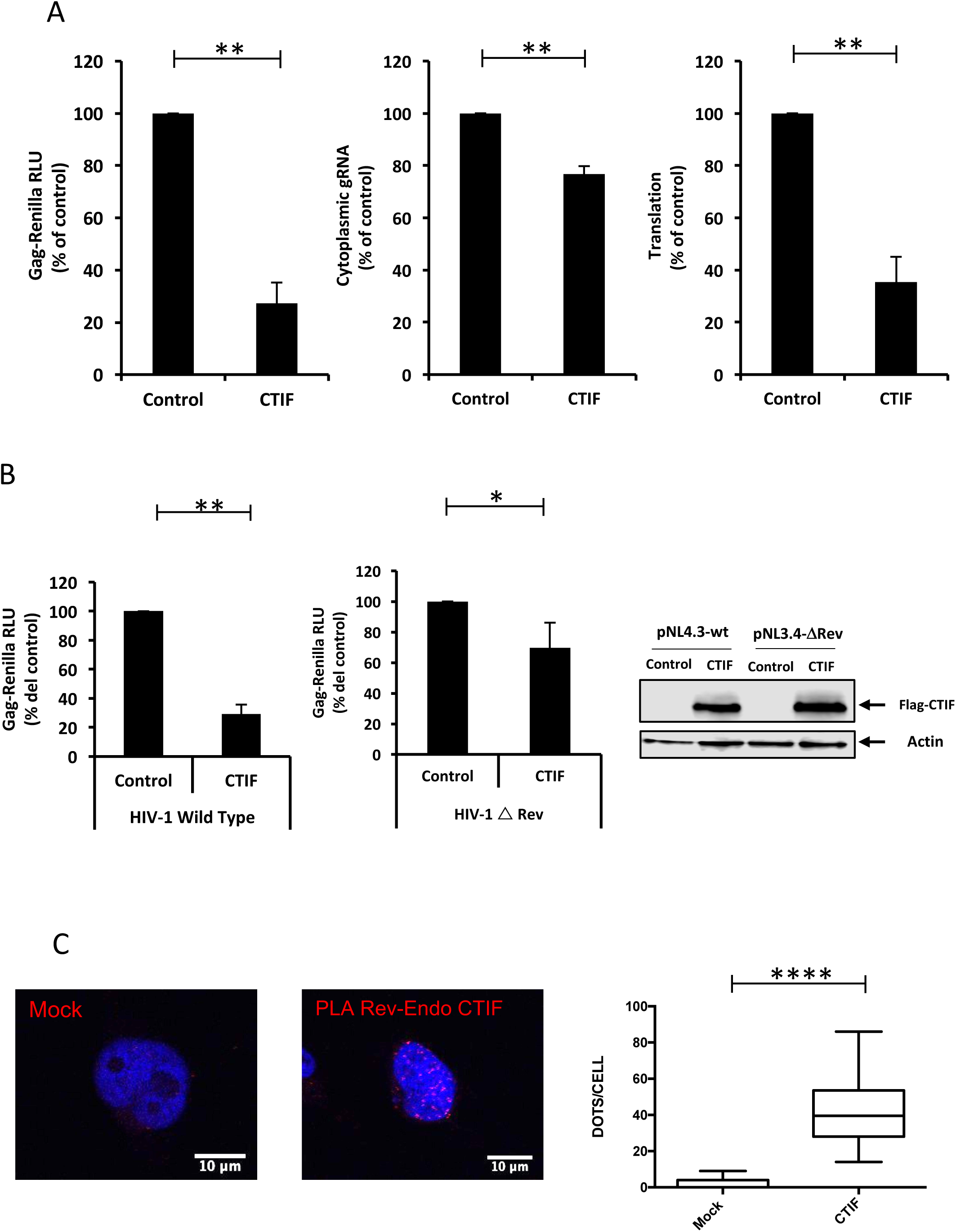
CTIF acts on Rev activity. A) HeLa cells were transfected with 1 µg of pCDNA-d2EGFP (used as a control) or pCDNA-Flag-CTIF together with 0,3 µg pNL4.3R and Renilla activity (left panel), cytoplasmic unspliced mRNA levels (middle panel) and unspliced mRNA translation (right panel) were determined 24 hpt. Results were normalized to the control (arbitrary set to 100%) and correspond to the mean +/- SD of three independent experiments (**P < 0.01, t-test). B) HeLa cells were transfected with 1 µg of pCDNA-d2EGFP (used as a control) or pCDNA-Flag-CTIF together with 0,3 µg of pNL4.3R or pNL4.3R-ΔRev and Renilla luciferase activity was determined at 24 hpt. Results were normalized to the control (arbitrary set to 100%) and correspond to the mean +/- SD of three independent experiments (*P < 0.05; **P < 0.01, t-test). Extracts from HeLa from different conditions were used to verify CTIF overexpression by Western blot. Actin were used as loading control. C) HeLa cells transfected with 1 µg pEGFP-Rev were subjected to the proximity ligation assay using a rabbit anti-CTIF antibody (rabbit) and a mouse anti-GFP antibody and the Duolink^®^ *in situ* kit as described in materials and methods. Mock corresponds to untransfected cells. Quantification of dots per cell in Mock-CTIF (n= 34 cells) and Rev-CTIF (n= 33 cells) are presented (****P < 0.0001, Mann–Whitney test).

Since the viral protein Rev is the major post-transcriptional regulator of the unspliced mRNA by allowing the efficient nuclear export and translation of the viral transcript, we then sought to determine whether Rev was involved in the inhibition of Gag expression mediated by CTIF. Thus, we analyzed the impact of Flag-CTIF expression on Gag synthesis in the presence or absence of Rev. Despite our ΔRev provirus produced lower amounts of Gag when compared to the wild type provirus (data not shown), we observed that CTIF was unable to exert the strong inhibition of Gag synthesis in the absence of Rev, suggesting that CTIF may at least target the function of Rev during unspliced mRNA translation (Fig. 2B). As such, our proximity ligation assay revealed that Flag-Rev interacts with endogenous CTIF (Fig. 2C and Supplementary Fig. 2).

Taking together, these data suggest that CTIF interacts with Rev and targets the function of the viral protein during translation of the unspliced mRNA.

### CTIF inhibits Gag synthesis through its N-terminal CBP80-binding domain

CTIF was shown to contain two major domains, the N-terminal region (CTIF 1-305) that contains the CBP80-binding domain and C-terminal region (CTIF 306-598), which harbors a MIF4G domain (25) (Fig. 3A). Thus, in order to determine whether the inhibitory activity of CTIF was exerted by one of these domains, we evaluated their impact on Gag synthesis from the pNL4.3R reporter provirus (Fig. 3B). We observed that ectopic expression of the N-terminal domain alone was able to inhibit the synthesis of Gag in a similar level compared with the full-length protein. The C-terminal domain was unable to interfere with Gag synthesis indicating that the N-terminal domain contains the inhibitory activity of CTIF. Similar results were obtained by Western blot using the pNL4.3 provirus and an anti-Cap24 antibody indicating that these effects are not exerted on the Renilla luciferase activity (Supplementary Fig. 3A).

**Figure 3:**
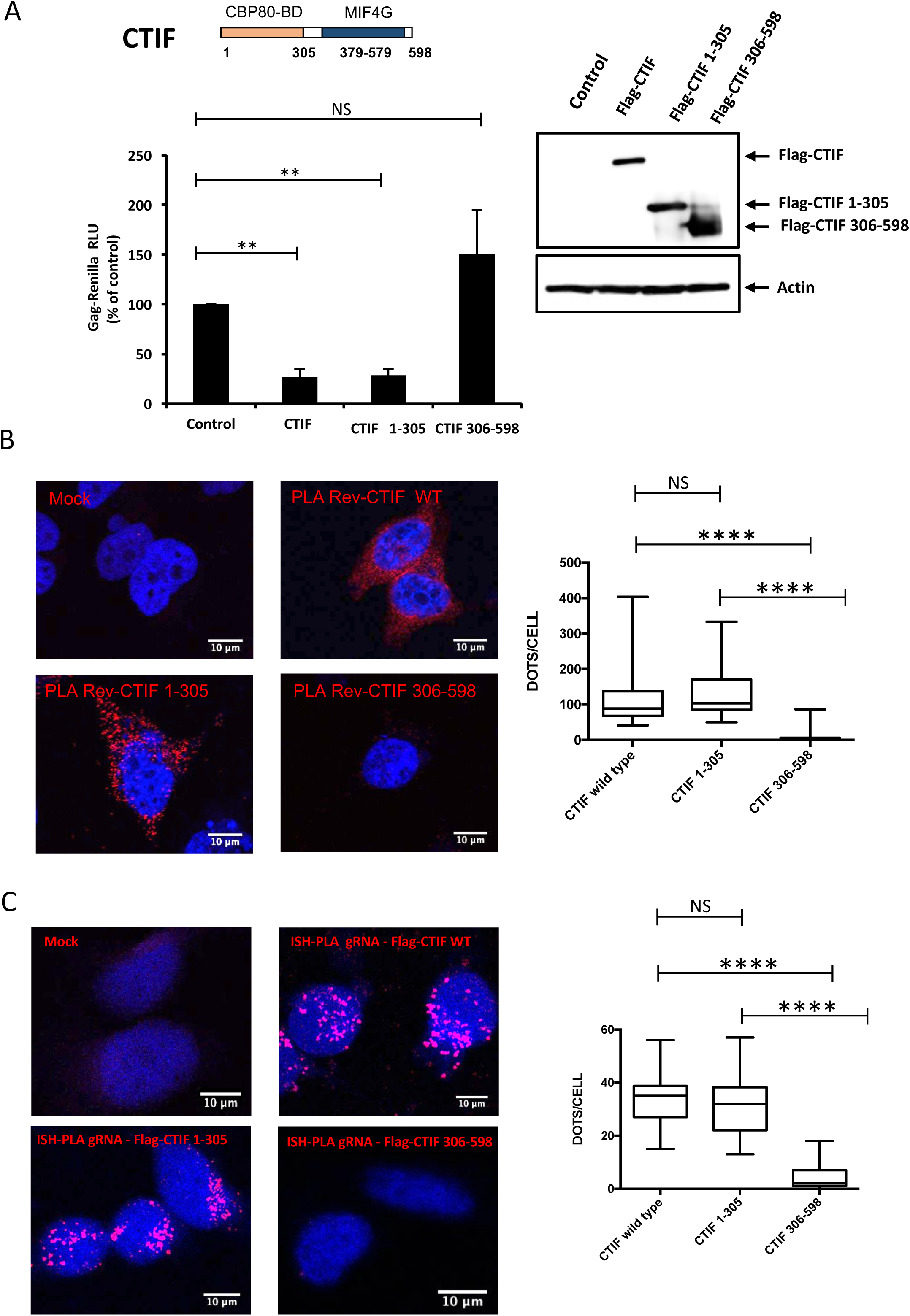
CTIF inhibits Gag synthesis through its N-terminal domain. A) Upper panel: Scheme of CTIF. The CBP80-binding domain and the MIF4G domain are indicated. Numbers indicate amino acid positions. Bottom panel: HeLa cells were transfected with 1 µg of pCDNA-d2EGFP (used as a control), pCDNA3-Flag-CTIF, pCDNA3-Flag-CTIF(1-305) or pCDNA-Flag-CTIF(306-598) together with 0,3 µg of pNL4.3R as described in materials and methods. Renilla activity was determined at 24 hpt. Results were normalized to the control (arbitrary set to 100%) and correspond to the mean +/- SD of three independent experiments (**P < 0.01; NS= non-significant, t-test). Right panel: In parallel, cells extracts were used to detect Flag-CTIF by Western blot for expression control. Actin were used as a loading control. HeLa cells transfected with 1 µg of pEGFP-Rev together with 1 µg of pCDNA3-Flag-CTIF, pCDNA-Flag-CTIF(1-305) or pCDNA-Flag-CTIF(306-598) were subjected to the proximity ligation assay using a rabbit anti-Flag antibody (rabbit) and a mouse anti-GFP antibody and the Duolink® *in situ* kit. Mock corresponds to untransfected cells. Quantification of dots per cell in Rev-CTIF (n = 32 cells), Rev-CTIF(1-305) (n = 44 cells) and Rev-CTIF(306-598) (n = 43 cells) are presented (****P < 0.0001, NS; non-significant, Mann–Whitney test). C) HeLa cells transfected with pNL4.3 together with pCDNA3-Flag-CTIF, pCDNA-Flag-CTIF(1-305) or pCDNA-Flag-CTIF(306-598) were subjected to ISH-PLA as described (Ref). Mock corresponds to untransfected cells. Quantification of dots per cell in unspliced mRNA-CTIF (n = 36 cells), unspliced mRNA-CTIF(1-305) (n = 30 cells) and unspliced mRNA-CTIF(306-598) (n = 27 cells) are presented (****P < 0.0001, NS; non-significant, Mann– Whitney test).

Consistent with the N-terminal domain as the responsible of the inhibitory effect of CTIF on Gag synthesis, we observed that this domain, but not the C-terminal domain, interacts with EGFP-Rev similar to the wild type protein (Fig. 3B). Interestingly, we observed that the Rev-CTIF interaction mainly occurs in the cytoplasm consistent with CTIF targeting the cytoplasmic functions of Rev on translation of the unspliced mRNA (see below).

We then used *in situ* hybridization coupled to the proximity ligation assay (ISH-PLA) in order to evaluate whether full-length CTIF and its isolated domains were recruited to the unspliced mRNA ribonucleoprotein complex during viral replication. We observed that both full-length CTIF and the N-terminal domain form complexes with the viral transcript (Fig. 3C). Consistent with its inability to interact with Rev and to interfere with Gag synthesis, we observed that the C-terminal domain of CTIF is not recruited to the unspliced mRNA. It should be mentioned that neither full-length CTIF nor the isolated domains altered the subcellular localization of the unspliced mRNA as judged by RNA FISH and confocal microscopy analyses (Supplementary Fig. 3B).

Taking together, these data suggest that CTIF interacts with Rev through its N-terminal domain and this interaction allows the recruitment of CTIF to the unspliced mRNA leading to the inhibition in Gag synthesis.

### CTIF affects the subcellular localization of Rev and its interaction with CBP80

From data presented above, it seems that CTIF and Rev mainly interacts in the cytoplasm despite most of the viral protein is normally found in the cell nucleus concentrated at the nucleolus. Thus, we sought to determine whether CTIF has an impact on the subcellular localization of EGFP-Rev. As expected, EGFP-Rev signal was mainly present in the nucleus/nucleolus of the cells under control conditions. However, we observed a significant increase of the cytoplasmic signal of EGFP-Rev when either full-length CTIF or the N-terminal domain were expressed together with the viral protein (Fig. 4A). Quantification of the subcellular localization of EGFP-Rev indicates that under control conditions near of the 90% of the cells exhibit an exclusive nuclear localization of EGFP-Rev (Fig. 4B). In contrast, we observed that 70-80% of the cells presented a cytoplasmic localization of EGFP-Rev when full-length CTIF or the N-terminal domain were co-expressed further confirming that the N-terminal domain of CTIF contains the inhibitory activity on Gag synthesis (Fig. 4B). Consistent with its inability to interact with Rev and interfere with Gag synthesis, the C-terminal domain of CTIF did not impact on the subcellular localization of Rev (Figs. 4A and 4B).

**Figure 4:**
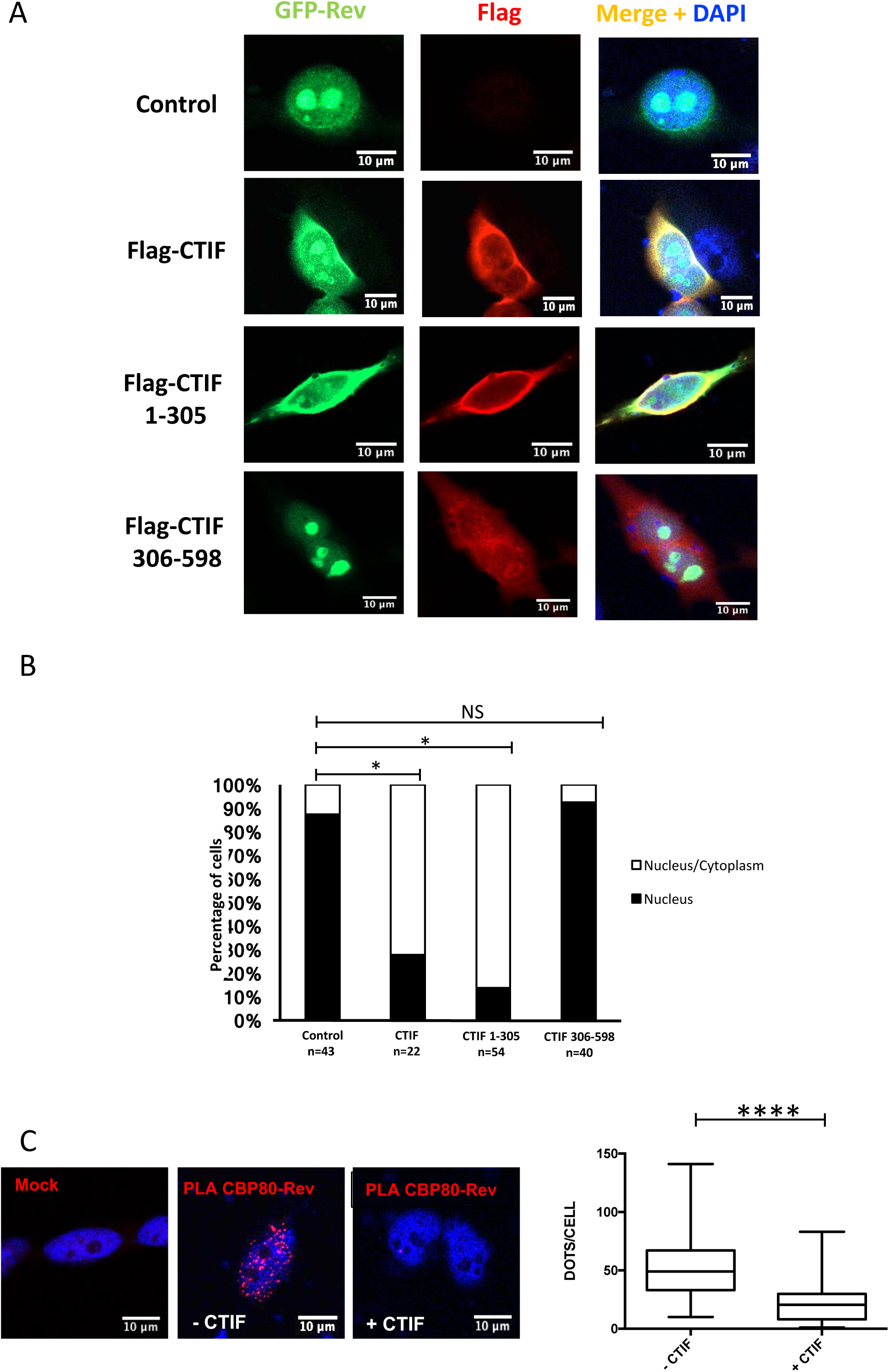
CTIF interferes with the Rev-CBP80 interaction. A) HeLa cells transfected with pEGFPC1-Rev together with pCDNA3-Flag-CTIF, pCDNA3-Flag-CTIF(1-305), pCDNA3-Flag-CTIF(306-598) or pCDNA3-Renilla (used as a control) were subjected to immunofluorescence assay using a rabbit anti-Flag antibody. Signals for DAPI (blue), EGFP-Rev (green), Flag (Red) and Merge are presented (scale bar 10 μm). B) Quantification of the sub-cellular localization of EGFP-Rev in conditions presented in A) (*P < 0.05; NS; non-significant, t-test). C) HeLa cells transfected with pEGFP-Rev and pCMV-Myc-CBP80 together with pCDNA3-Renilla (- CTIF condition) or pCDNA3-Flag-CTIF (+ CTIF condition) were analyzed for the Rev-CBP80 interaction by PLA as described in materials and methods. Mock corresponds to untransfected cells. (B) Quantification of dots per cell in – CTIF (n = 43 cells) and + CTIF (n = 37 cells) is presented (****: P < 0.0001, Mann–Whitney test).

We recently reported that Rev associates with CBP80 to promote nuclear export and translation of the unspliced mRNA. Thus, we wanted to investigate whether the inhibitory activity of full-length CTIF was related with this interaction. Quantification of our PLA experiments showed that expression of CTIF strongly interferes with the interaction between Rev and CBP80 (Fig. 4C and Supplementary Fig. 4).

Taking together, these data suggest that CTIF induces a cytoplasmic retention of Rev and interferes with the association between the viral protein and CBP80. This lack of interaction between Rev and CBP80 in the presence of CTIF may result in the inhibition of Gag synthesis.

### CTIF-mediated restriction is conserved in human lentiviruses

We finally wanted to determine whether the negative effect of CTIF was exclusive for Rev-containing human lentiviruses or was conserved in simple retroviruses or other RNA viruses. For this, we first evaluated the effect of CTIF expression on an HIV-2 reporter provirus. We observed that similar to HIV-1, Gag synthesis from the HIV-2 unspliced mRNA was also inhibited by CTIF (Fig. 5A). This result correlate with the presence of the regulatory protein Rev in both complex human lentiviruses. Thus, in order to corroborate whether the presence of Rev is determinant for the inhibition mediated by CTIF, we evaluated the effect of this cellular protein on Gag synthesis from the simple retrovirus Murine Leukemia Virus (MLV), which do not express a Rev-like regulatory protein and also on human Respiratory Syncytial Virus (hRSV), which is a non-related negative stranded RNA virus (Figs. 5B and 5C). We analyzed the effects of CTIF overexpression on the synthesis of Gag from MLV and the fusion protein (F-protein) from hRSV but none of these viruses was affected, indicating that CTIF is not a pan-antiviral protein but rather selectively targets human lentiviruses expressing the viral protein Rev.

**Figure 5:**
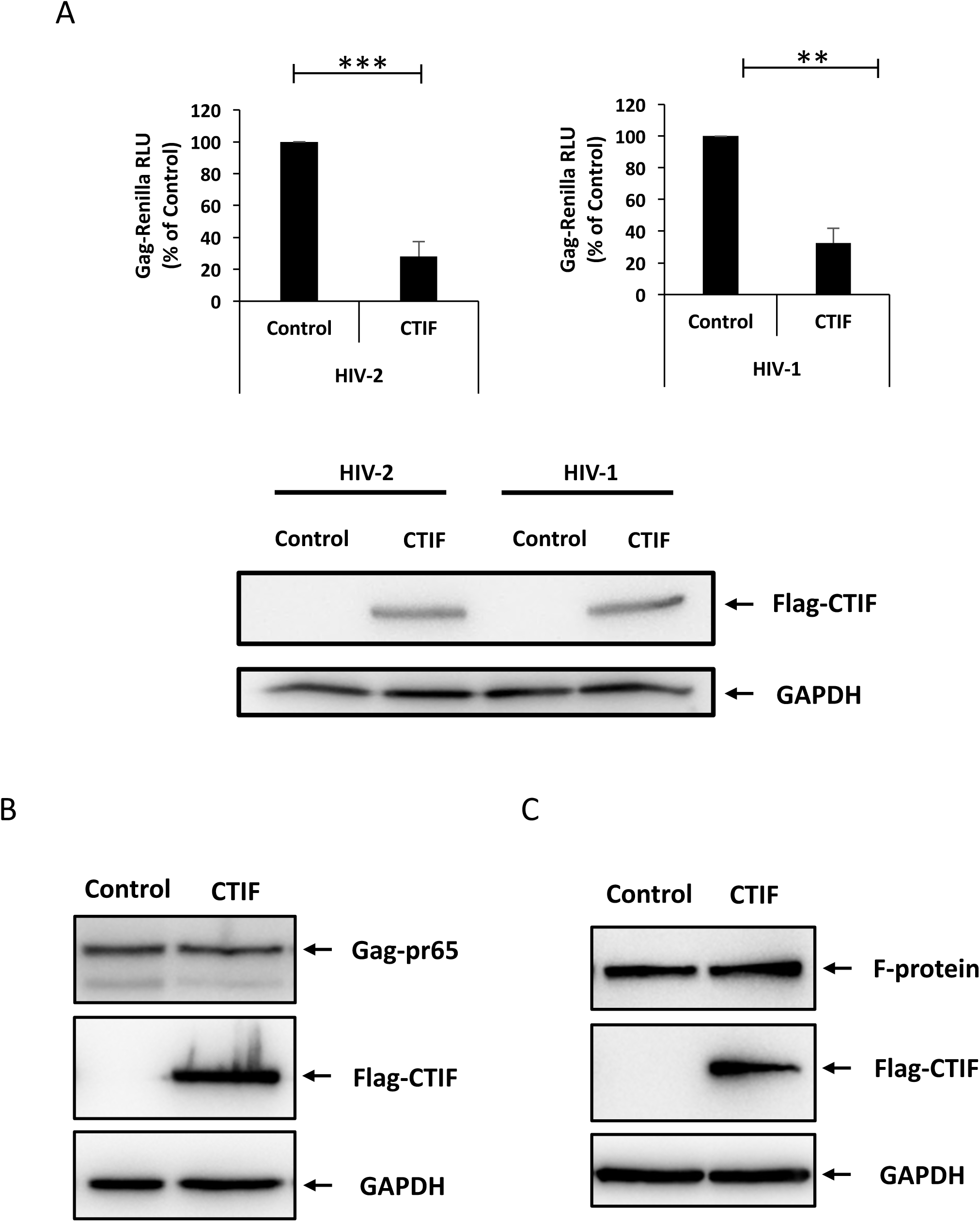
CTIF restricts Rev expressing human lentiviruses. A) HeLa cells were transfected with 1 µg of pcDNA-d2EGFP (used as control) or pcDNA3-Flag-CTIF together with 0.3 µg of pNL4.3R (HIV-1) or pROD10R (HIV-2) and Renilla activity was determined at 24 hpt. Results were normalized to the control (arbitrary set to 100%) and correspond to the mean +/- SD of three independent experiments (**P < 0.01, ***P < 0.001; t-test). In parallel, cells extracts were used to detect Flag-CTIF by Western blot for expression control. Actin were used as a loading control. B) HEK293T cells were transfected with 1.5 µg of pcDNA-d2EGFP (Control) or pcDNA3-Flag-CTIF together with 0.5 µg of pNCAC. Cells extract were prepared 24 hpt and subjected to Western blot to detect CAp30 of MLV, Flag-CTIF. GAPDH was used as a loading control. C) HEK293T cells were transfected with 1 µg of pcDNA-d2EGFP (Control) or pcDNA3-Flag-CTIF and 6 h post-transfection the cells were infected with hRSV MOI: 1. Cells extracts were prepared at 24 hpi and subjected to Western blot to detect the viral F protein and Flag-CTIF. GAPDH was used as a loading control.

## DISCUSSION

From an RNA biology point of view, Gag synthesis from the HIV-1 unspliced mRNA should not be as efficient as it is. This is because this viral transcript possesses several features known to be incompatible with efficient nuclear export and translation in mammalian cells. First, it contains functional splice donors and acceptors sites and thus, it needs to subvert the cellular splicing machinery in order to accumulate in the nucleus of infected cells. Second, the lack of intron removal avoids the splicing-dependent recruitment of nuclear host factors and thus, the unspliced mRNA is not assembled into a canonical mRNP that exits the nucleus associated to NXF1 and is directed to the translational machinery in the cytoplasm. Third, it harbors a highly structured 5’-UTR expected to interfere with the cap-dependent ribosomal scanning mechanism of translation initiation. Despite all these functional constraints, the HIV-1 unspliced mRNA reaches the cytoplasm efficiently and is highly specialized in ribosome recruitment producing huge amounts of Gag protein during viral replication. The viral protein Rev orchestrates the post-transcriptional regulation of the unspliced mRNA by acting as a multifunctional bridge between host factors and the viral transcript. By binding the Rev response element (RRE) present at the 3’-UTR of the unspliced mRNA through an RNA binding domain and the host karyopherin CRM1 through its nuclear export signal (NES), Rev ensures the cytoplasmic accumulation of the viral mRNA. Rev has also been involved in ribosome recruitment of the unspliced mRNA although the mechanisms at play are poorly understood. Recent work including ours reported that Rev interacts with the CBC subunit CBP80. In our study, we demonstrated that CBP80 cooperates with Rev during nuclear export and translation of the unspliced mRNA. We also showed that the unspliced mRNA preferentially associates with CBP80 and that Rev promotes this association. Since we reported that Rev favored the association of a complex also containing the DEAD-box RNA helicase eIF4AI but not other translation initiation factors such as eIF4GI or eIF3g, we decided to further investigate the composition of this unusual viral mRNP. We focused on CTIF since it is the translation initiation factor that scaffolds the CBC-bound mRNP and the 40S ribosomal subunit through interactions with CBP80 and eIF3g. However and to most of our surprise, we observed that CTIF rather inhibits HIV-1 replication by blocking Gag synthesis. An interesting point of our observations is that CTIF levels increased at early hours post-infection to then decrease when Gag started to accumulate suggesting that CTIF would not be necessary or deleterious for the synthesis of this viral protein. Since CTIF seems not to be an interferon stimulated gene (33), we believe that endogenous CTIF levels must be increased by the virus in order to promote the accumulation of early viral gene products such as Tat and Rev, which are expressed from multiply spliced transcripts and follows a canonical gene expression pathway. Then, HIV-1 may induce the decrease in the levels of CTIF in order to accumulate the structural protein Gag. In support of this, we showed that low level of CTIF promoted the accumulation of Gag, while high levels of CTIF were inhibitory for Gag synthesis. Indeed, CTIF binds Rev through its N-terminal domain inducing the cytoplasmic accumulation of the viral protein and impeding the assembly or inducing the disassembly of the Rev-CBP80 complex (Fig. 6).

**Figure 6:**
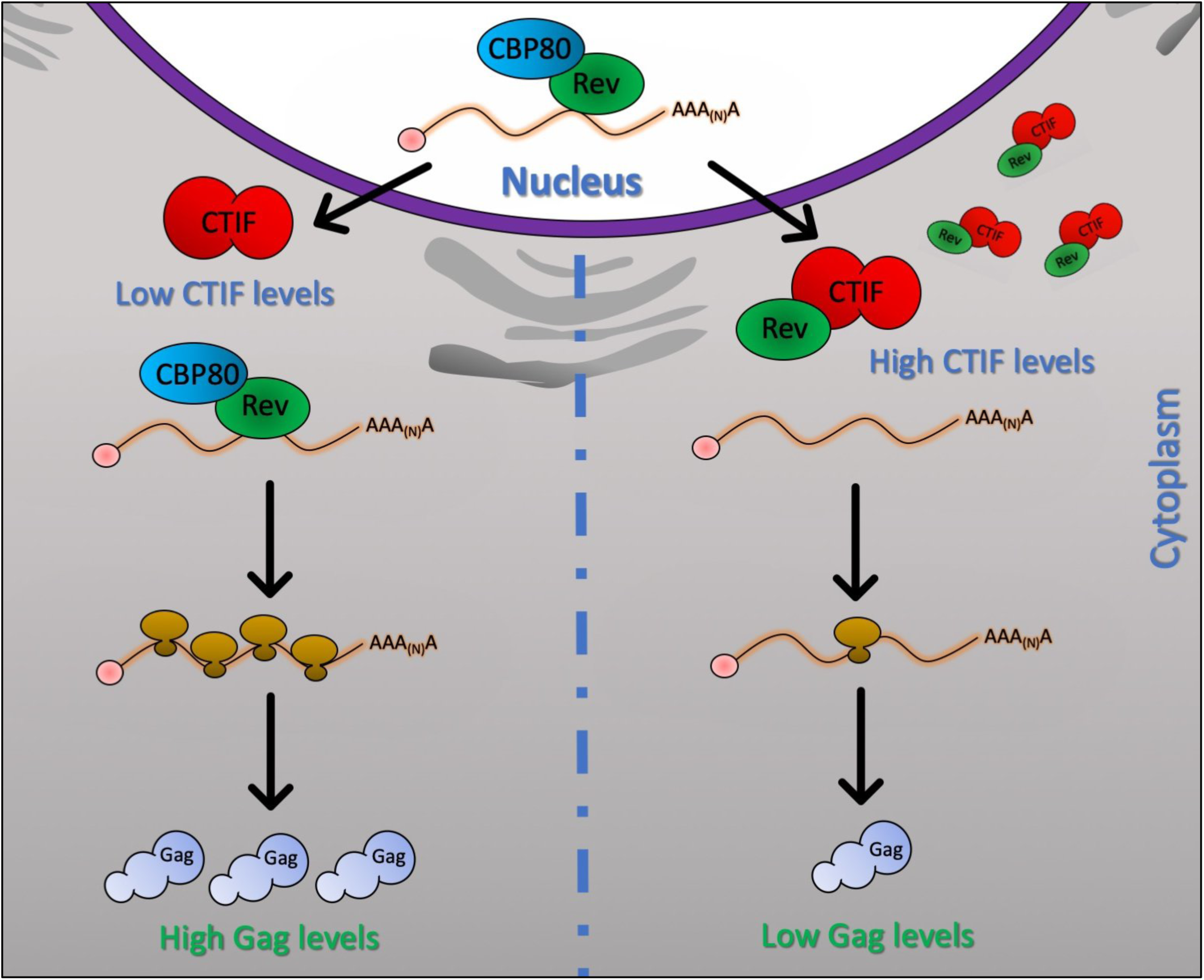
A model for the activity of CTIF on HIV-1 Gag synthesis. The HIV-1 unspliced mRNA associates with Rev and CBP80 in the nucleus and this complex is exported to the cytoplasm. Under high levels of CTIF, CBP80 is displaced from Rev and the CTIF-Rev complex is inefficient in supporting unspliced mRNA translation resulting in low levels of Gag synthesis (left). When CTIF levels decrease, high levels of Gag synthesis are ensured by the loading of the CBP80-Rev complex onto de unspliced mRNA (right).

Considering the different origins of HIV-1 and HIV-2, the specific inhibition of Gag synthesis observed in both human lentiviruses and the high conservation of CTIF between human and non-human primates (data not shown), it is tempting to speculate that the inhibition of replication mediated by CTIF must be conserved at least in primate lentiviruses.

The small molecule ABX464 was shown to interfere with HIV-1 replication by targeting the function of Rev and CBP80 and is currently in a phase II clinical trial. This information together with our results confirm the critical relevance of the Rev-CBP80 complex on HIV-1 replication and highlights the usefulness of this interaction as a reliable target for the development of novel antiretroviral therapies aimed to interfere with viral gene expression from the unspliced mRNA.

## ACKNOWLEDGEMENTS

Authors are especially grateful of Dr. Yoon Ki Kim (Korea University, Korea) for critical reading of the manuscript and his helpful comments as well as for kindly sharing all the CTIF-expressing plasmids used in this study. Authors wish also thank to Dr. Gloria Arriagada (Universidad Andrés Bello, Chile) for sharing the anti-MLV CA antibody. The following reagents were obtained through the NIH AIDS Reagents Program, Division of AIDS, NIAID, NIH: HIV-1 p24 Monoclonal Antibody (183-H12-5C) from Dr. Bruce Chesebro and Kathy Wehrly.

---

**Supplementary Fig. 1.**
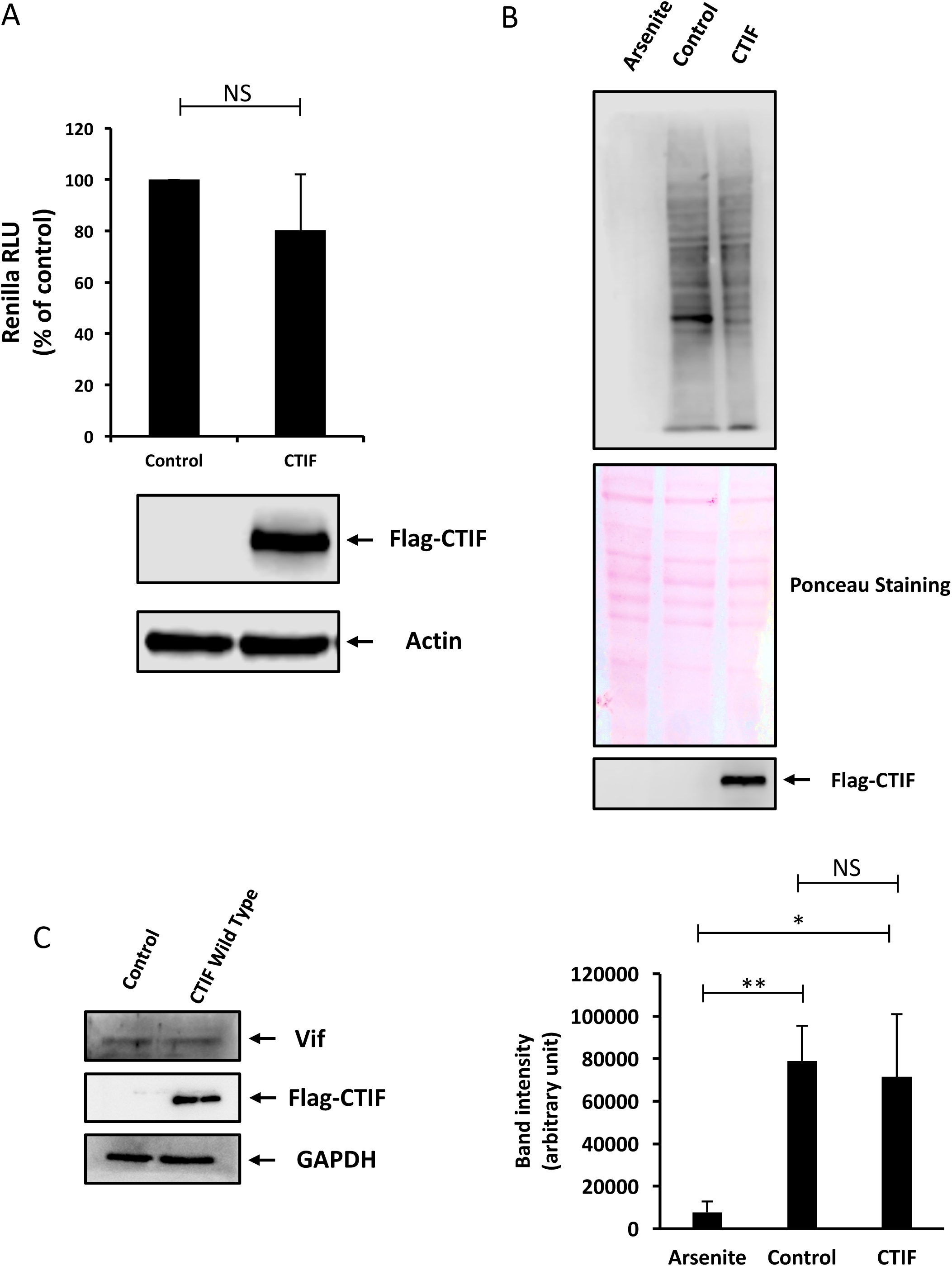
(A) HeLa cells were transfected with 0.3 µg of pcDNA-d2EGFP (used as a control) or pcDNA3-Flag-CTIF together with 1µg of pCIneo-Renilla as described in materials and methods. Renilla activity were determined 24 hpt. Results were normalized to the control (arbitrary set to 100%) and correspond to the mean +/- SD of three independent experiments. (NS; non-significant, t-test). In parallel, cells extracts were used to detect Flag and GFP by Western blot for expression control. Actin were used as a loading control. (B) Upper panel: HeLa cells were transfected with 1 µg of pcDNAd2EGFP (Control) or pcDNA3-Flag-CTIF and treated with puromycin as described in materials and methods. Cell extract were used to detect puromycin by Western blot. Ponceau staining were used for loading control. Cells extracts were used to detect Flag-CTIF by Western blot for expression control. GFP band were signaling with an arrow. Arsenite were used for positive inhibition protein synthesis control. Bottom panel: Band intensity quantification of puromycin Western blot and correspond to the mean +/- SD of three independent experiments (*P < 0.05; **P < 0.01 and NS; non-significant, t-test). (C) HeLa cells were transfected with pcDNAd2EGFP (Control), pcDNA3-Flag-CTIF, together with pNL4.3 as described in materials and methods. Cells extract were used to detect Vif by Western blot. We detect Flag and GFP by Western blot for expression control. GAPDH were used as a loading control.

**Supplementary Fig. 2.**
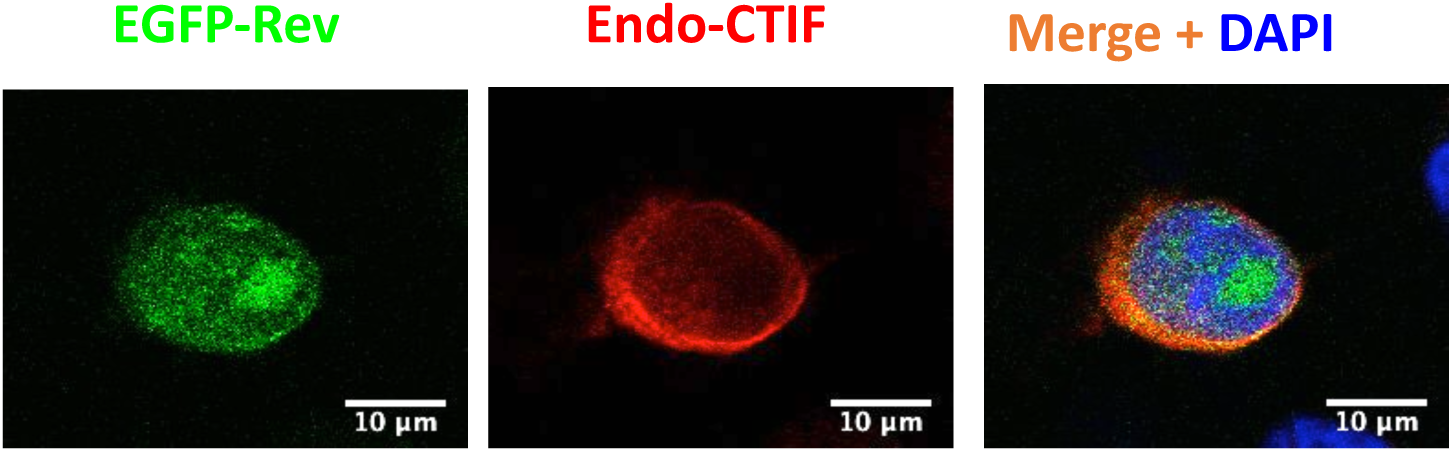
HeLa cells transfected with 1µg pEGFP-Rev were subjected to immunofluorescence as described in materials and methods. EGFP-Rev is shown in green and endogenous CTIF in red. Scale bar 10 µm.

**Supplementary Fig. 3.**
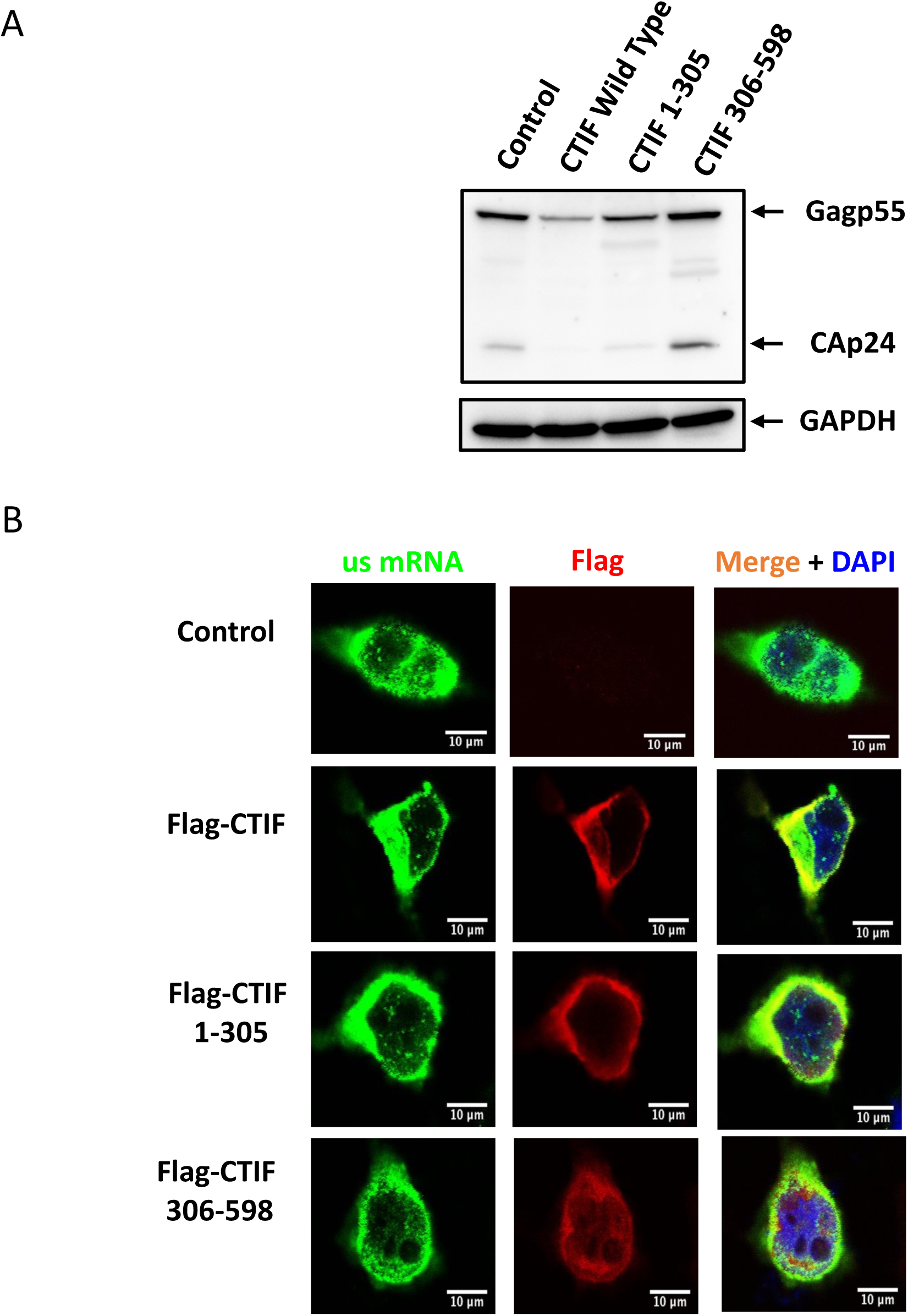
A) HeLa cells were transfected with pcDNAd2EGFP (Control), pcDNA3-Flag-CTIF, pcDNA3-Flag-CTIF(1-305) or pcDNA-Flag-CTIF(306-598) together with pNL4.3 as described in materials and methods. Cells extract were used to detect CAp24 by Western blot. GAPDH were used as a loading control. B) HeLa cells transfected with 1µg of pNL4.3 together with 1µg of pUC19 (used as a control), pcDNA3-Flag-CTIF, pcDNA3-Flag-CTIF(1-305) or pcDNA3-Flag-CTIF(306-598) were subjected to FISH as described in materials and methods. The unspliced mRNA is shown in green and Flag-CTIF in red. Scale bar 10 m.

**Supplementary Fig. 4.**
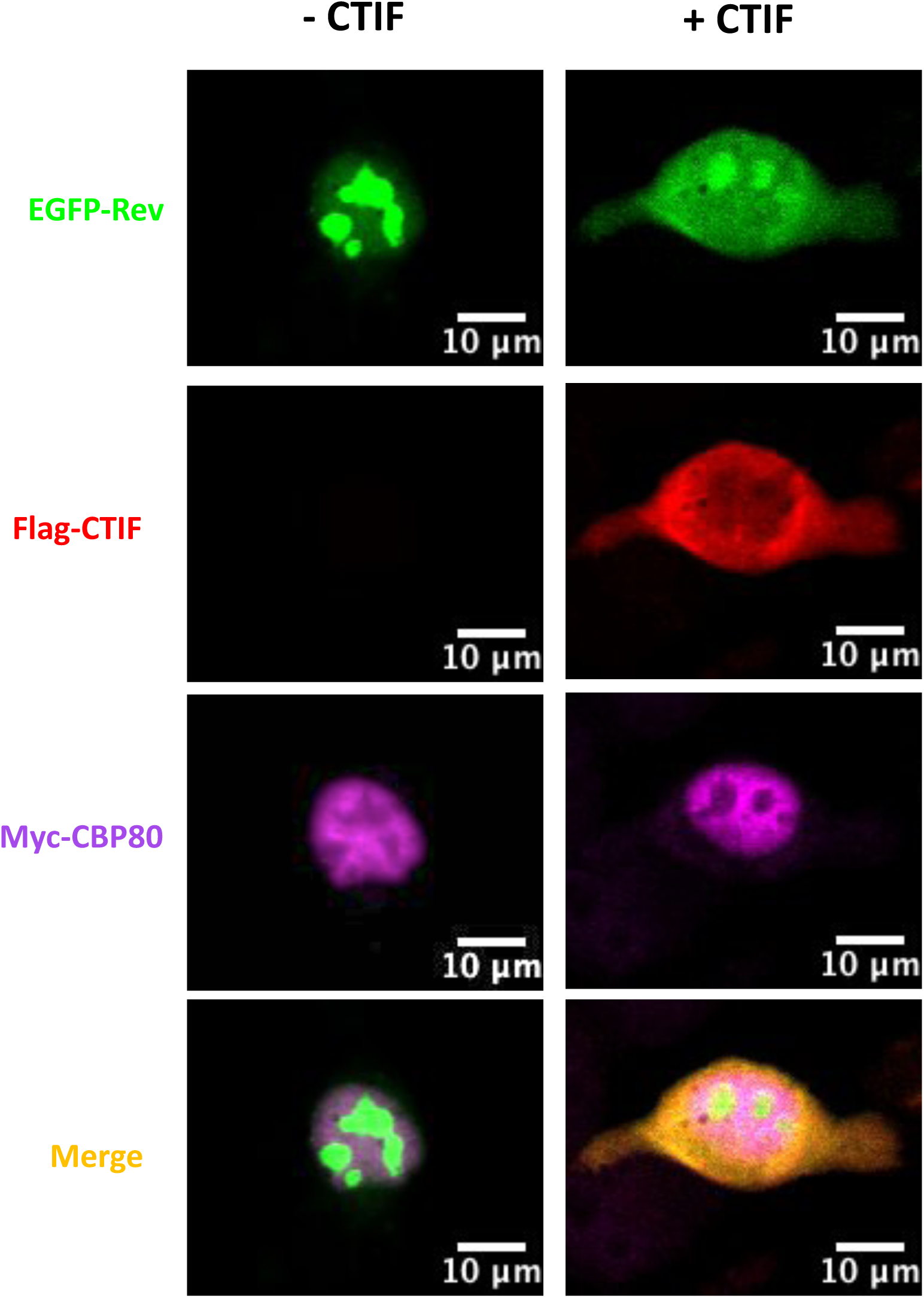
HeLa cells transfected with pEGFP-Rev and pCMV-Myc-CBP80 together with pCDNA3-Renilla (- CTIF condition) or pCDNA3-Flag-CTIF (+ CTIF condition) were were subjected to IF as described in materials and methods. The GFP-Rev is shown in green, Flag-CTIF in red and Myc-CBP80 in magenta. Scale bar 10 m.

**Supplementary Table 1:**
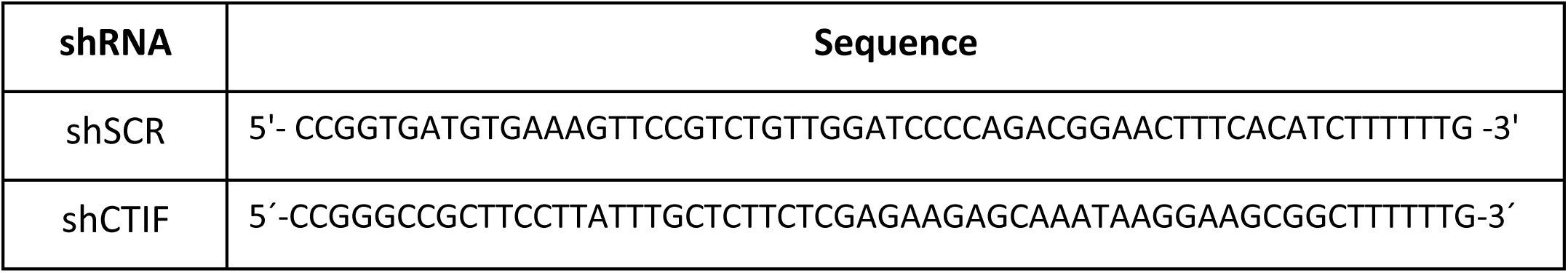
sequences of the shRNA used in this study

**Supplementary Table 2:**
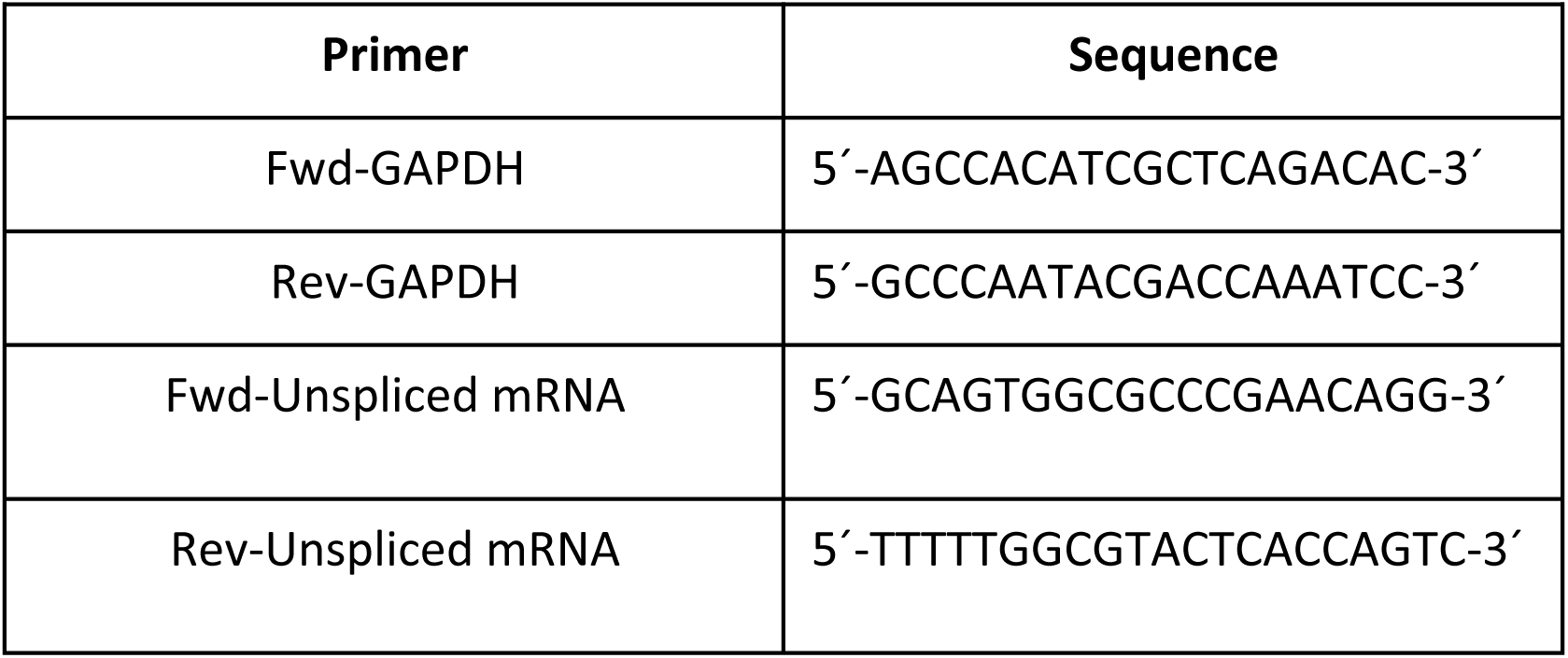
Primers used in RT-qPCR assays

